# Insight on Selective Breeding the Robustness Based on Field Survival Records: New Genetic Evaluation of Survival Traits in White-leg Shrimp *(Penaeus vannamei)* Breeding Line

**DOI:** 10.1101/2022.08.14.503924

**Authors:** Shengjie Ren, Peter B. Mather, Binguo Tang, David A. Hurwood

**Author notes:** **Correspondence:** Shengjie Ren.

## Abstract

Survival is an old trait in animal breeding, yet commonly neglected nowadays for its simple binary records and low levels of heritability in aquaculture species. These traits however, can provide valuable field data when selecting for robustness in genetic improvement programs. In the current study, linear multivariate animal model (LMA) was used for the genetic analysis of survival records from two-year classes (BL2019 and BL2020) of white-leg shrimp (*Penaeus vannamei*) breeding lines with a total number of 52, 248 individuals from 481 fullsib families recorded for data collection. During grow-out test period, 10 day intervals of survival data were considered as independent traits. Two survival definitions, binary survivability (S) and continuous survival in days (SL), were used for the genetic analysis of survival records to investigate; i) whether adding more survival time information could improve estimation of genetic parameters; ii) the trajectory of survival heritability across time, and iii) patterns of genetic correlations of survival traits across time. Levels of heritability estimates for both S and SL were low (0.005 to 0.076), while heritability for survival day number was found to be similar with that of binary records at each observation time and were highly genetically correlated (*r*_*g*_ >0.8). Heritability estimates of body weight (BW) for BL2019 and BL2020 were 0.486 and 0.373, respectively. Trajectories of survival heritability across time showed a gradual increase across the grow-out test period but slowed or reached a plateau during the later grow-out test period. Genetic correlations among survival traits in the grow-out tests were moderate to high, and the closer the times were between estimates, the higher were their genetic correlations. In contrast, genetic correlations between both survival traits and body weight were low but positive. Here we provide the first report of the trajectory of heritability estimates for survival traits across time in aquaculture. Results will be useful for developing robust improved white-leg shrimp culture strains in selective breeding programs based on field survival data.

## 1 INTRODUCTION

Aquaculture is playing an increasingly important role in world food security, development of economic sustainability, and to provide practical solutions for addressing ecosystem services issues (Houston et al., 2020; Naylor et al., 2021; Bernatchez et al., 2017). Sustainability of aquaculture production in global food systems however, has become vulnerable due to the rapid expansion of aquaculture industry, outbreak of diseases/pathogens, water environmental pollution, and in particular, from the impacts of climate changes (Reid et al., 2019; Troell et al., 2014). Consequently, there has seen an increasing demand for better management and breeding of robust culture lines (Friggens et al., 2017). Genetic improvement via selective breeding is widely acknowledged as an efficient tool to improve economically important traits including: feed conversion ratios, biomass production, and overall survival rates of domesticated aquatic animals (Hung et al., 2013; Gjedrem et al., 2012; Gjedrem and Rye, 2018; Nguyen, 2016). A wide range of projects have confirmed selective breeding to be an efficient strategy for enhancing overall survival rate and disease resistance performance in farmed aquatic species, and thereby contributing to the development of aquaculture in a sustainable way (Houston, 2017; Yáñez et al., 2014).

In general terms, two selection approaches have been commonly used to develop robustness in aquaculture genetic breeding programs. One approach is to improve specific disease resistance via controlled challenge tests in target breeding lines. For this approach, tested families are artificially infected with a specific pathogen in a controlled environment condition via intra-peritoneal injection, immersion or cohabitation. Following this, additive genetic variance among families for phenotypic resistance against the target pathogen can be identified and used in future breeding plans (Ødegård et al., 2011; Yáñez et al., 2014). Currently, disease resistance strains have been developed successfully via this approach for a number of aquatic species, including farmed salmonid species (Correa et al., 2015; Vallejo et al., 2017; Barría et al., 2019), Pacific oyster (Gutierrez et al., 2018), and European sea bass (Palaiokostas et al., 2018). A second approach is to use selection based on survival data records in the field and this can provide another important data source for selecting robustness by improving overall individual survival rate (Gjedrem and Rye, 2018) and for developing specific disease resistance strains (Barría et al., 2020; Fraslin et al., 2022). This approach allows direct collection of data under real commercial farm conditions, and thereby avoids potential for genotype-by-environment (G-by-E) problems as seen in controlled challenge experiments between challenge test environments and production conditions on farm. Survival data records however, are often neglected in aquaculture breeding programs because they are based on simple binary records (0 or 1) and generally show low heritability.

Genetic analyses of survival phenotypic data in aquaculture breeding programs are commonly treated as a binary trait with ‘alive vs dead’ reported for individuals at a specific observation time point scored as 1 and 0, respectively. Under this scenario, individuals that died early or later during the grow-out period would be given the same score, which means that useful information about relative survival time and/or lifespan is lost and therefore has not been used in the genetic analysis (Ødegård et al., 2006). Moreover, binary record variables of survival are commonly non-normally distributed, an issue that may compromise estimations of genetic components using a linear mixed animal model. To address this problem, survival records scored as continuous traits of survival time have been implemented successfully for genetic evaluation of survival data in animal breeding programs (Ducroq and Casella, 1996; Van Pelt et al., 2015; Heise et al., 2016). In this way, survival phenotype can be recorded as normally distributed continuous traits and information about individuals with different survival time/lifespans can be captured effectively in the analysis. To date, genetic analyses of survival data scored as continuous traits have only been reported on farmed aquatic species for controlled challenge tests (Suebsong et al., 2019; Joshi et al., 2021a) while survival data from commercial breeding lines has rarely been investigated.

Different genetic evaluation models are available for genetic analyses of survival traits in animal breeding (Forabosco et al., 2009). A proportional hazard model (PHM) is a common model used for genetic evaluation of survival traits, and can be applied with a free open access software package in *Survival Kit* (Ducrocq and Sölkner, 1994; Ducrocq et al., 2010). PHM can handle large survival data sets rapidly and easily, manage survival data sets with skewed distributions, and is considered to provide accurate estimates (Sewalem et al., 2010; Zavadilová et al., 2011). In genetic evaluations however, it is difficult to process genetic correlations with other continuous traits at the same time (Tarrés et al., 2006). Alternatively, linear models and threshold models have been widely used for genetic analyses on survival data in aquaculture genetics. Compared with linear models, threshold models applying a logit link function are feasible for dealing with non-normally distributed binary survival data. However, they generally require more computational time and cannot estimate genetic correlations with other continuous traits simultaneously. In practice, analyses using either threshold models or PHM take almost five to ten times more computational time than do applying linear models to address almost the same tasks (Boettcher et al., 1999). Moreover, estimations of true breeding values (EBVs) have shown very similar correlations between linear models and threshold models (Veerkamp et al., 2002). While different linear models including random regression models (RRMs) have been used successfully for national genetic evaluations of survival data in dairy cattle (Sasaki et al., 2015; Van Pelt et al., 2015, 2016; Heise et al., 2016, 2018), there are currently no standardized model choices for routine genetic evaluation of survival data in aquaculture breeding programs.

Another important consideration for genetic analysis of survival traits in aquaculture species is changes with time. Survival traits are dynamic quantitative traits, which can change spatially and temporally due to multiple interactions between animal and environmental constraints. Therefore, understanding the trajectory of time for survival traits can be very useful for making critical decisions in a breeding plan (Schaeffer, 2004). Studies of trajectory of time for important economic traits in aquaculture genetics are still rare, and the few reported cases have mostly focused on growth traits (Turra et al., 2012; He et al., 2018; Schlicht et al., 2018), while to date, there have been no reports on trajectory of survival traits across time.

White-leg shrimp (*Penaeus vannamei*) has become the most widely farmed prawn species across the world. Annual global production reached ∼4.4 million tons with a commercial value of 26.7 billion USD in 2020, which ranked as the most important traded food commodity across the aquaculture sector (Kumar et al., 2016; FAO, 2020). Sustainability of prawn farming has been affected however, by emergence of several diseases that show high mortality rates (Trang et al., 2019). Developing robust culture strains of white-leg shrimp via selective breeding will play a crucial role for improving the economic profitability and animal welfare of this major aquatic farmed species. The main objective here was therefore to develop routine genetic evaluation of survival records in a genetic improvement program for white-leg shrimp in China. Specifically, we evaluated: (1) genetic parameters from two different survival definitions for the binary traits of survivability and the continuous trait of survival time; (2) the trajectory of heritability for survival traits across the grow-out test period; and (3) genetic correlation patterns for survival traits across times and their correlation with growth. Results from the current program will provide insight on selection breeding of robust culture strains based on field survival records in aquaculture species.

## 2 MATERIALS AND METHODS

### Study Population

The study population constituted the breeding nucleus from a white-leg shrimp stock improvement program in Hainan Island (China), with the selection target being for local farm environments in China based on a family selection approach. Foundation populations were produced in 2015 sourced from 12 hatchery lines in China representing four genetic populations as evidenced from a population structure analysis (Ren et al., 2018). In 2015, 98 full families were produced (Ren et al., 2020a). Following this, for each breeding cycle, 209-250 families were produced over a one-week period. Family pedigree management used physical visible implant elastomer (VIE) tags, while in parallel a parentage assignment panel (Ren et al., 2022) was developed in 2019 to meet the demands of larger family numbers in the selective breeding program. The mating system for this program used a nested mating design via one single male with two females. The ratio of dam/sire was maintained at ∼ 1.7 with the aim to generate better genetic tier for EBVs estimation. Grow-out management conditions of the breeding line have been reported in earlier studies (Ren et al., 2020b, c).

### Data Records

In the current study, genetic analysis data were sourced from two different year classes (2019 (BL2019) and 2020 (BL2020)) in the breeding nucleus line. The BL2019 line, consisted of 243 full-sibs families generated from 157 sires and 243 dams over a one-week period while the BL2020 line consisted of 238 full-sibs families produced by 143 sires and 238 dams (**Table 1**). The mean number of shrimp per family for the grow-out test was 102.8 for BL2019 and 114.7 for BL2020, with a total number of 52, 248 individuals recorded for data collection (**Table 1**).

**Table 1.** Data structure of white-leg shrimp breeding nucleus lines.

| Year | No. shrimp | Shrimp/family | No. family | Sires | Dams |
| --- | --- | --- | --- | --- | --- |
| 2019 | 24, 980 | 102.8 | 243 | 157 | 243 |
| 2020 | 27, 304 | 114.7 | 238 | 143 | 238 |
| Total/mean | 52, 284 | 108.7 | 481 | 300 | 481 |

Dead individuals from the breeding lines were collected three times per day across the grow-out period and pedigree information of mortalities was recorded from visible implant elastomer (VIE) tags. At the end of the grow-out test stage, all remaining harvested individuals in the test system were scored for body weight, pedigree of family ID, and gender.

### Trait Definition

Data collected on survival from each 10 day interval were considered as independent traits over the grow-out period. Survival phenotype was recorded using two definitions in the genetic analysis. First, survival phenotype at the end of each 10 day period was coded as a binary trait of survivability, with 0 for dead individuals and 1 for live individuals. Secondly, survival data for each 10 day period was considered to be a continuous trait of survival time indicating how many days individuals survived over the test period. For example, live individuals in each breeding line at surviving to 100 days were coded as 100d and dead individuals at 86 days or 56 days were coded as 86d or 56d, respectively. Therefore, survival data records from BL2019 breeding line across the 117 days grow-out period were coded as 12 survivability binary traits (S1, S2, …, S11, S12) and 12 continuous traits of survival (SL1, SL2, …, SL11, SL12) for each observed time window of 10 days. Similarly, the 98 days records of survival data from BL2020 considered 10 survival binary traits (S1, S2, …, S9, S10) and 10 continuous traits of survival days (SL1, SL2, …, SL9, SL10), respectively. At the conclusion of the grow-out tests, phenotype of body weight for each shrimp was collected as a continuous trait (BW), while data from the final population census of live shrimps were recorded as a binary survival trait (SUR).

### Statistical Analysis

#### Survival Probability Estimates and Daily Mortality

Changes in survival probability of shrimp in breeding lines across the experimental grow-out period were analysed using the Kaplan-Meier estimator (Kaplan and Meier, 1958). Shrimp that survived at each observed time (days) were treated as censored. The “Survminer” function (Kassambara et al., 2017) in the **R** package (R Core Team,2019) was used to estimate Kaplan-Meier survival curves by;

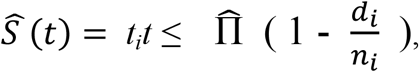

where, *n*_*i*_ is the number of alive shrimp at risk at observed time *t*_*i*_, and *d*_*i*_ is the number of deaths at the observed time. In addition, daily mortality in the breeding lines was also plotted for the general husbandry management assessment purposes.

#### Genetic Analyses Genetic parameters

WOMBAT software (Meyer, 2007) was used to fit the following linear multivariate animal model (LMA) for the genetic analysis:

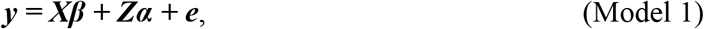

where, ***y*** is a vector of phenotype for the traits defined here (S1-S12, SL1-SL12, SUR, and BW); ***β*** is the vector of fixed effects including sex, tanks and family batches; ***α*** is the vector of random additive genetic effects; ***e*** is the vector of random residual errors; and, ***X*** and ***Z*** are known incidence matrices relating observations to the fixed and random effects mentioned above. Both vector of ***a*** and ***e*** are assumed to be multivariate normal distribution with mean zero and variances as:

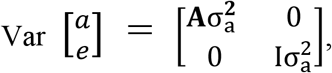

here, 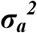 and 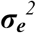 are the random additive variances and error variances, respectively. **A** is the numerator relationship matrix based on pedigree information, and **I** represents an identity matrix. Total variance 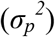 was calculated as the sum of random additive genetic variance 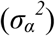 and random residual components 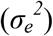. Heritability (*h*^*2*^) was calculated as the ratio of the random genetic variance to the total phenotype variance, 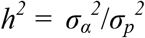. Phenotypic correlation (*r*_*p*_) between two traits was calculated as:

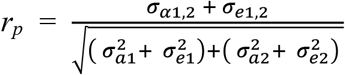 and genetic correlation (*r*_*g*_) between two traits was calculated as: 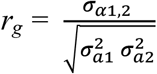. The values of data matrix for *r*_*p*_ and *r*_*g*_ were displayed as heat maps in **R** package (R Core Team,2019). From the output of LMA genetic analyses, *r*_*p*_ and *r*_*g*_ between phenotype traits were aligned using **R** package in the ‘Matrix’ package and after, function of ‘pheatmap’ in R was used to plot the graphic representations of the correlation matrix data.

#### Correlations of binary traits and survival days

Bivariate analyses were performed, in which the two definition types of survival phenotype were modeled simultaneously. Equations used in the bivariate animal model can be defined as follows:

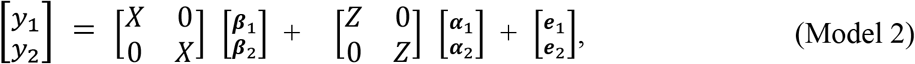

where, the symbols represent the same vectors as described in the multivariate analysis of Model 1; the subscripts 1 and 2 are two different records for survival data of binary traits (S1-S12) and survival days (SL1-SL12) at each observed time window, respectively.

#### Common environmental effect

Two linear univariate animal models were developed for the significance test of the full-sib family effect (*c*^*2*^) in the genetic analyses. Equations of univariate animal models can be written as follows:

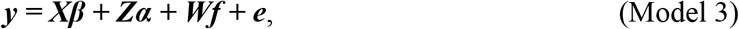

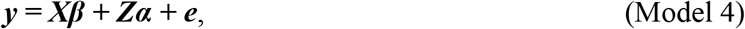

where, **f** is the vector of random full-sib family effects and **W** is the corresponding design matrices; other symbols represent the same vectors as described in Model 1. Compared with Model 3, there was no random full-sib family effect in Model 4. Likely statistical significance for the random full-sib family effects were analysed using a likelihood ratio approach. After running the two above univariate Models, significance for the random full-sib family effects were compared to the final log-likelihood (Maximum log L) using a Chi-square (*χ*^*2*^) test.

## 3 RESULTS

### Patterns of Survival

Kaplan–Meier survival curves showed changes in survival probability across the grow-out test periods for each year breeding line over the two year test period (**Figure 1**). At the end of the grow-out test period, survival probability for BL2019 was 89.4%, while for BL2020 it was 84.4%. These survival estimators however, presented some bias to the final population census (80.4% and 64.2%) at the end of grow-out test. Differences between the bias estimates indicate a proportion of shrimp mortality events apparently were not detected over the test periods. There were also significant differences (*P*<0.01) in mortality among families for the two breeding lines investigated here. Means of daily mortality records for BL2019 and BL2020 were 0.1% and 0.17%, respectively (**Figure 1**). Overall, daily mortalities fluctuated at 0.1%-0.25% for breeding lines across the two years. Small peaks in mortality were first observed at the beginning of each test in the first week, which in part, can be explained as due to handling stress during VIE tagging. Most daily mortality peaks were evident at later stages of the grow-out test periods, where the highest daily mortality at 0.38% peaked at ∼ 90 days for BL2019, while the highest daily mortality for BL2020 peaked at 0.78% at 70 days. The time period of the highest daily mortality events coincided with the coldest winter season period at the hatchery location in Hainan.

**Figure 1.**
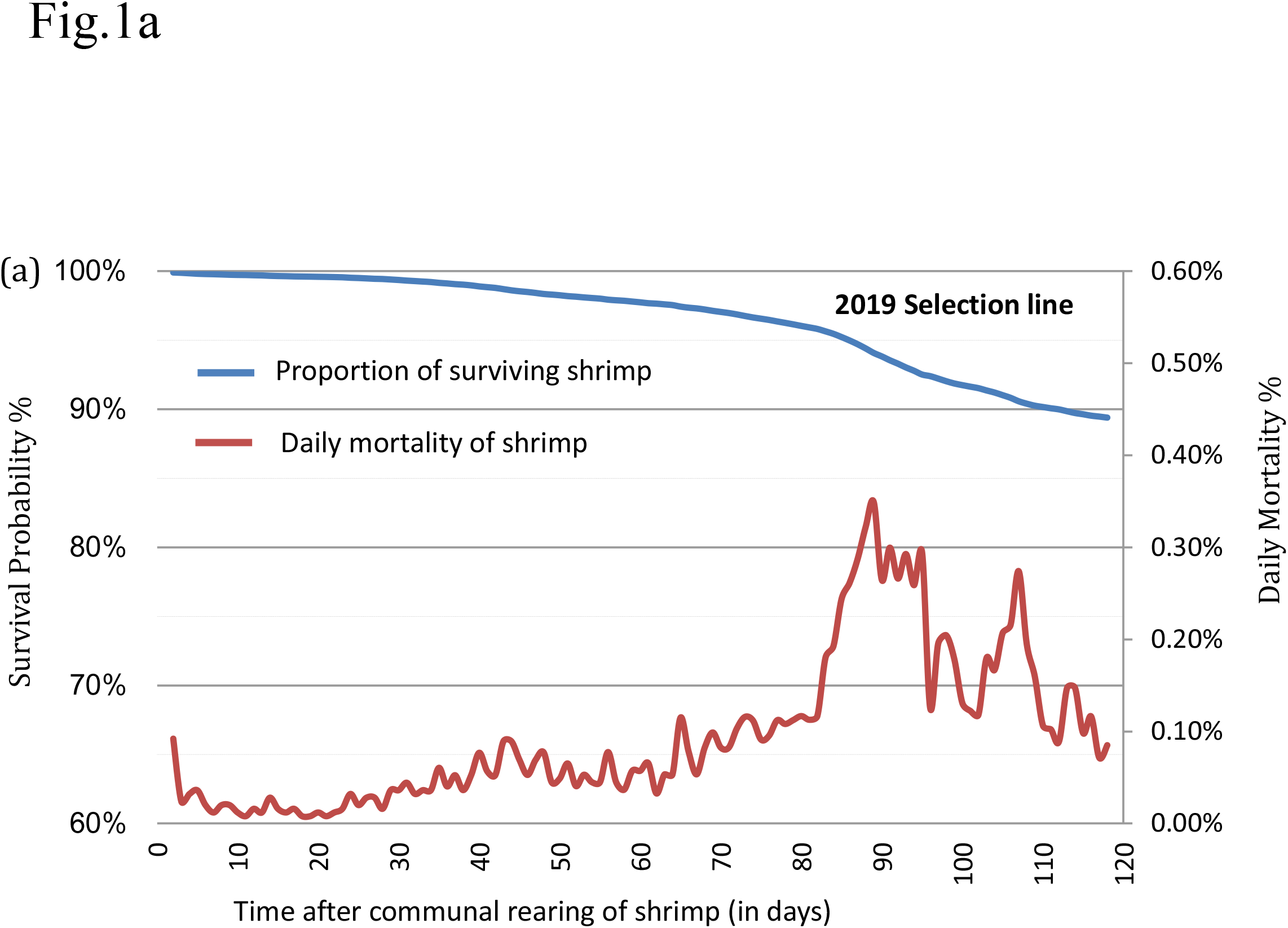
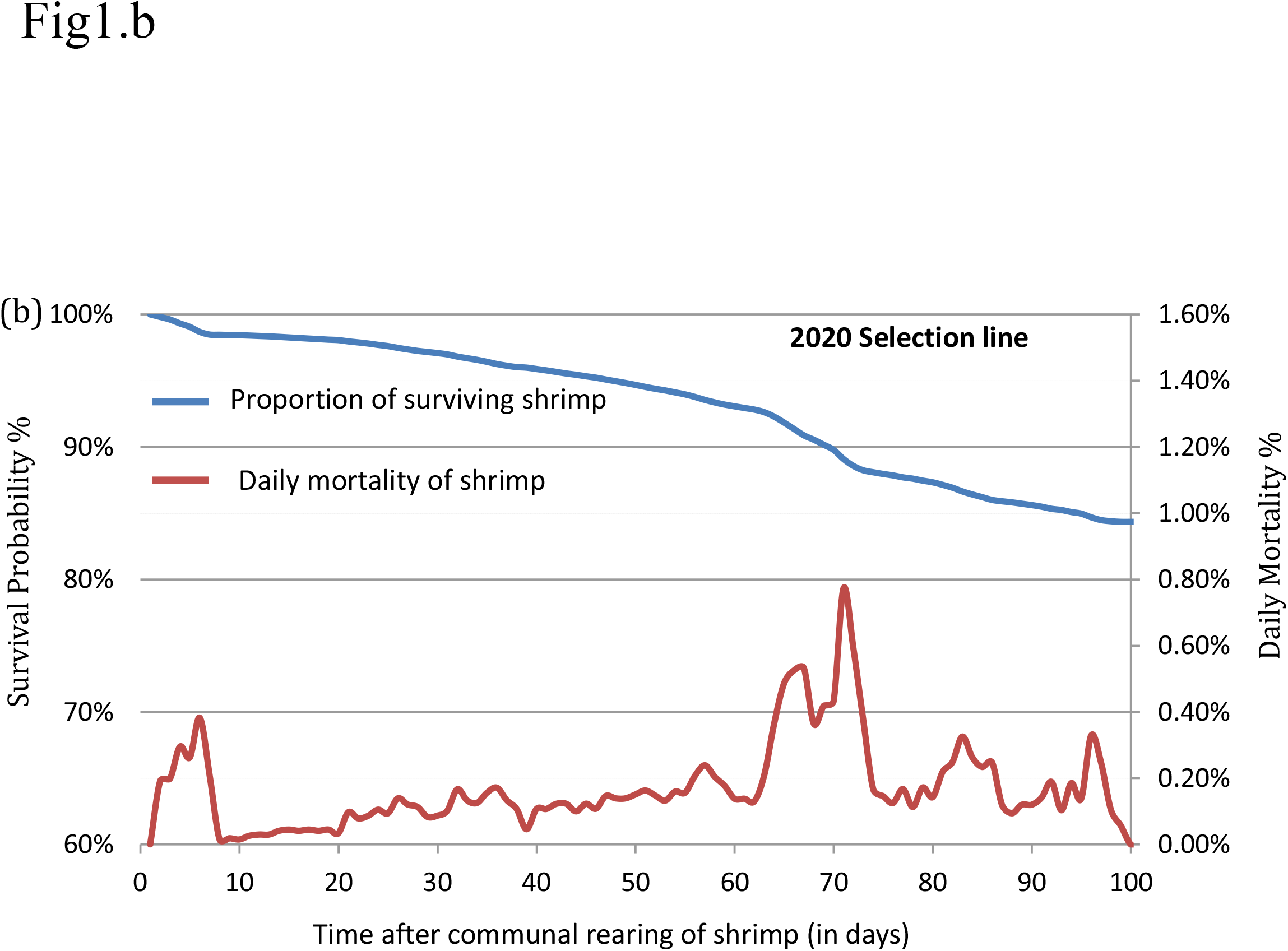
Kaplan-Meier survival curves and daily mortality changes for the shrimp breeding lines: (**a**) 2019 (BL2019) of 243 fullsib families and (**b**) 2020 (BL2020) of 238 fullsib families.

### Descriptive Statistics of Genetic Analyses

For both year breeding lines, all families had records for survival data. Basic descriptive statistics for each trait are presented in **Table 2**. The coefficient of variation (CV) for survival traits gradually increased for both two types of survival data record (S and SL), in both lines. CV for binary survival traits ranged from 4.82% to 32.79% for BL2019, while levels of CV for survival days were much lower compared with binary traits. Similar patterns for CV were also observed for the BL 2020 line.

**Table 2.** Descriptive statistics of phenotype traits for the breeding lines of BL2019 and BL2020.

| Trait | *BL2019* |  |  |  |  | *BL2020* |  |  |  |  |
| --- | --- | --- | --- | --- | --- | --- | --- | --- | --- | --- |
|  | *N* | Min | Max | Mean | CV% | *N* | Min | Max | Mean | CV% |
| S1 | 24980 | 0.000 | 1.000 | 0.998 | 4.824 | 27304 | 0.000 | 1.000 | 0.984 | 12.576 |
| S2 | 24980 | 0.000 | 1.000 | 0.996 | 6.211 | 27304 | 0.000 | 1.000 | 0.981 | 13.914 |
| S3 | 24980 | 0.000 | 1.000 | 0.994 | 7.927 | 27304 | 0.000 | 1.000 | 0.972 | 16.872 |
| S4 | 24980 | 0.000 | 1.000 | 0.990 | 10.095 | 27304 | 0.000 | 1.000 | 0.959 | 20.724 |
| S5 | 24980 | 0.000 | 1.000 | 0.984 | 12.786 | 27304 | 0.000 | 1.000 | 0.947 | 23.759 |
| S6 | 24980 | 0.000 | 1.000 | 0.979 | 14.551 | 27304 | 0.000 | 1.000 | 0.932 | 26.931 |
| S7 | 24980 | 0.000 | 1.000 | 0.973 | 16.682 | 27304 | 0.000 | 1.000 | 0.899 | 33.482 |
| S8 | 24980 | 0.000 | 1.000 | 0.963 | 19.471 | 27304 | 0.000 | 1.000 | 0.876 | 37.557 |
| S9 | 24980 | 0.000 | 1.000 | 0.942 | 24.724 | 27304 | 0.000 | 1.000 | 0.865 | 39.538 |
| S10 | 24980 | 0.000 | 1.000 | 0.922 | 29.025 | 27304 | 0.000 | 1.000 | 0.859 | 40.512 |
| S11 | 24980 | 0.000 | 1.000 | 0.910 | 31.370 | NA | NA | NA | NA | NA |
| S12 | 24980 | 0.000 | 1.000 | 0.903 | 32.794 | NA | NA | NA | NA | NA |
| SL1 | 24980 | 1.000 | 10.000 | 9.986 | 3.320 | 27304 | 1.000 | 10.000 | 9.904 | 8.037 |
| SL2 | 24980 | 1.000 | 20.000 | 19.957 | 4.063 | 27304 | 1.000 | 20.000 | 19.730 | 10.350 |
| SL3 | 24980 | 1.000 | 30.000 | 29.909 | 4.680 | 27304 | 1.000 | 30.000 | 29.493 | 11.549 |
| SL4 | 24980 | 1.000 | 40.000 | 39.833 | 5.305 | 27304 | 1.000 | 40.000 | 39.134 | 12.680 |
| SL5 | 24980 | 1.000 | 50.000 | 49.704 | 6.013 | 27304 | 1.000 | 50.000 | 48.657 | 13.856 |
| SL6 | 24980 | 1.000 | 60.000 | 59.521 | 6.818 | 27304 | 1.000 | 60.000 | 58.045 | 15.037 |
| SL7 | 24980 | 1.000 | 70.000 | 69.288 | 7.638 | 27304 | 1.000 | 70.000 | 67.215 | 16.234 |
| SL8 | 24980 | 1.000 | 80.000 | 78.978 | 8.458 | 27304 | 1.000 | 80.000 | 76.058 | 17.555 |
| SL9 | 24980 | 1.000 | 90.000 | 88.535 | 9.317 | 27304 | 1.000 | 90.000 | 84.753 | 18.953 |
| SL10 | 24980 | 1.000 | 100.000 | 97.862 | 10.286 | 27304 | 1.000 | 98.000 | 93.352 | 20.332 |
| SL11 | 24980 | 1.000 | 110.000 | 107.024 | 11.364 | NA | NA | NA | NA | NA |
| SL12 | 24980 | 1.000 | 118.000 | 116.089 | 12.489 | NA | NA | NA | NA | NA |
| SUR | 24980 | 0.000 | 1.000 | 0.755 | 56.923 | 27304 | 0.000 | 1.000 | 0.585 | 90.720 |
| BW | 20078 | 2.300 | 57.000 | 21.394 | 26.891 | 17529 | 1.900 | 41.900 | 17.946 | 23.626 |

*N*, the number of shrimp; S1-S12, survivability of binary traits for each observed time window of 10 days period; SL1-SL12, continuous traits of survival days for each observed time window of 10 days period; SUR, binary trait of survival for the final population census; BW, body weight of shrimp.

Means of survival trait (S) at the end of the grow-out test period were higher than that of SUR, indicating some mortality events had not been observed.

### Genetic Parameters

Estimates of variance components and heritability of survivability records for binary traits (S), survival days (SL), final population census of survival (SUR), and body weight (BW) are presented in **Table 3**. Levels of heritability estimates for S and SL were both low and ranged from 0.005 to 0.076. Heritability for survival days (SL) was found to be closer with that of binary records (S) at each time of observation. While slightly higher heritability estimates were found for S than for SL at early stages of the grow-out testing period in the BL2019 line, they were very closer in the middle to later test periods. The trajectories of S and SL during grow-out test period are illustrated in **Figure 2**. The slightly higher heritability estimates for survival in BL2020 than in BL2019 were expected since we recorded a higher daily mortality in BL2020. It was interesting that, for both two-year data groups, heritability of S showed a steady increase over time. While SL estimates increased during the first 60 days, levels of heritability for SL fluctuated at ∼0.06 and maintained a plateau after this initial period.

**Table 3.** Estimates of variance components and heritabilities (*h*^*2*^ *± se*) for the genetic analyse of survival data and body weight.

| Trait | *BL2019* |  |  |  | *BL2020* |  |  |  |
| --- | --- | --- | --- | --- | --- | --- | --- | --- |
| | $\sigma_a^2$ | $\sigma_p^2$ | $\sigma_e^2$ | $h^2 \pm se$ | $\sigma_a^2$ | $\sigma_p^2$ | $\sigma_e^2$ | $h^2$ |
| S1 | 1.09 E<sup>-5</sup> | 2.31 E<sup>-3</sup> | 2.30 E<sup>-3</sup> | 0.005 ± 0.002 | 3.02 E<sup>-4</sup> | 1.53 E<sup>-2</sup> | 1.50 E<sup>-2</sup> | 0.020 ± 0.004 |
| S2 | 6.64 E<sup>-5</sup> | 3.83 E<sup>-3</sup> | 3.76 E<sup>-3</sup> | 0.017 ± 0.003 | 5.88 E<sup>-4</sup> | 1.86 E<sup>-2</sup> | 1.80 E<sup>-2</sup> | 0.017 ± 0.003 |
| S3 | 2.11 E<sup>-4</sup> | 6.21 E<sup>-3</sup> | 6.00 E<sup>-3</sup> | 0.034 ± 0.005 | 1.16 E<sup>-3</sup> | 2.70 E<sup>-2</sup> | 2.58 E<sup>-2</sup> | 0.043 ± 0.006 |
| S4 | 4.59 E<sup>-4</sup> | 9.99 E<sup>-3</sup> | 9.53 E<sup>-3</sup> | 0.046 ± 0.005 | 2.20 E<sup>-3</sup> | 3.96 E<sup>-2</sup> | 3.74 E<sup>-2</sup> | 0.056 ± 0.007 |
| S5 | 9.57 E<sup>-4</sup> | 1.58 E<sup>-2</sup> | 1.49 E<sup>-2</sup> | 0.060 ± 0.007 | 3.11 E<sup>-3</sup> | 5.08 E<sup>-2</sup> | 4.77 E<sup>-2</sup> | 0.061 ± 0.007 |
| S6 | 1.27 E<sup>-3</sup> | 2.03 E<sup>-2</sup> | 1.90 E<sup>-2</sup> | 0.063 ± 0.008 | 4.38 E<sup>-3</sup> | 6.32 E<sup>-2</sup> | 5.89 E<sup>-2</sup> | 0.069 ± 0.008 |
| S7 | 1.45 E<sup>-3</sup> | 2.62 E<sup>-2</sup> | 2.48 E<sup>-2</sup> | 0.055 ± 0.007 | 6.31 E<sup>-3</sup> | 9.06 E<sup>-2</sup> | 8.43 E<sup>-2</sup> | 0.070 ± 0.008 |
| S8 | 2.08 E<sup>-3</sup> | 3.49 E<sup>-2</sup> | 3.29 E<sup>-2</sup> | 0.060 ± 0.007 | 8.21 E<sup>-3</sup> | 1.09 E<sup>-1</sup> | 1.00 E<sup>-1</sup> | 0.076 ± 0.009 |
| S9 | 3.16 E<sup>-3</sup> | 5.38 E<sup>-2</sup> | 5.07 E<sup>-2</sup> | 0.059 ± 0.007 | 8.88 E<sup>-3</sup> | 1.17 E<sup>-1</sup> | 1.08 E<sup>-1</sup> | 0.076 ± 0.009 |
| S10 | 4.85 E<sup>-3</sup> | 7.11 E<sup>-2</sup> | 6.62 E<sup>-2</sup> | 0.068 ± 0.008 | 9.15 E<sup>-3</sup> | 1.21 E<sup>-1</sup> | 1.12 E<sup>-1</sup> | 0.075 ± 0.009 |
| S11 | 5.32 E<sup>-3</sup> | 8.08 E<sup>-2</sup> | 7.55 E<sup>-2</sup> | 0.066 ± 0.008 | NA | NA | NA | NA |
| S12 | 5.86 E<sup>-3</sup> | 8.70 E<sup>-2</sup> | 8.12 E<sup>-2</sup> | 0.067 ± 0.008 | NA | NA | NA | NA |
| SL1 | 1.14 E<sup>-4</sup> | 1.10 E<sup>-1</sup> | 1.10 E<sup>-1</sup> | 0.001 ± 0.002 | 1.09 E<sup>-2</sup> | 6.34 E<sup>-1</sup> | 6.23 E<sup>-1</sup> | 0.017 ± 0.003 |
| SL2 | 3.79 E<sup>-3</sup> | 6.56 E<sup>-1</sup> | 6.52 E<sup>-1</sup> | 0.006 ± 0.002 | 9.25 E<sup>-2</sup> | 4.17 E<sup>0</sup> | 4.08 E<sup>0</sup> | 0.022 ± 0.004 |
| SL3 | 2.58 E<sup>-2</sup> | 1.96 E<sup>0</sup> | 1.93 E<sup>0</sup> | 0.013 ± 0.003 | 3.29 E<sup>-1</sup> | 1.16 E<sup>1</sup> | 1.13 E<sup>1</sup> | 0.029 ± 0.005 |
| SL4 | 1.11 E<sup>-1</sup> | 4.46 E<sup>0</sup> | 4.35 E<sup>0</sup> | 0.025 ± 0.004 | 9.16 E<sup>-1</sup> | 2.46 E<sup>1</sup> | 2.37 E<sup>1</sup> | 0.037 ± 0.005 |
| SL5 | 3.41 E<sup>-1</sup> | 8.93 E<sup>0</sup> | 8.58 E<sup>0</sup> | 0.038 ± 0.005 | 2.06 E<sup>0</sup> | 4.55 E<sup>1</sup> | 4.35 E<sup>1</sup> | 0.045 ± 0.006 |
| SL6 | 8.28 E<sup>-1</sup> | 1.65 E<sup>1</sup> | 1.56 E<sup>1</sup> | 0.050 ± 0.007 | 4.01 E<sup>0</sup> | 7.63 E<sup>1</sup> | 7.23 E<sup>1</sup> | 0.052 ± 0.007 |
| SL7 | 1.61 E<sup>0</sup> | 2.80 E<sup>1</sup> | 2.63 E<sup>1</sup> | 0.058 ± 0.007 | 7.05 E<sup>0</sup> | 1.19 E<sup>2</sup> | 1.12 E<sup>2</sup> | 0.059 ± 0.007 |
| SL8 | 2.80 E<sup>0</sup> | 4.45 E<sup>1</sup> | 4.17 E<sup>1</sup> | 0.063 ± 0.008 | 1.16 E<sup>1</sup> | 1.79 E<sup>2</sup> | 1.67 E<sup>2</sup> | 0.065 ± 0.008 |
| SL9 | 4.50 E<sup>0</sup> | 6.78 E<sup>1</sup> | 6.33 E<sup>1</sup> | 0.066 ± 0.008 | 1.79 E<sup>1</sup> | 2.59 E<sup>2</sup> | 2.41 E<sup>2</sup> | 0.069 ± 0.008 |
| SL10 | 6.98 E<sup>0</sup> | 1.01 E<sup>2</sup> | 9.37 E<sup>1</sup> | 0.069 ± 0.008 | 2.60 E<sup>1</sup> | 3.61 E<sup>2</sup> | 3.35 E<sup>2</sup> | 0.072 ± 0.008 |
| SL11 | 1.05 E<sup>1</sup> | 1.47 E<sup>2</sup> | 1.36 E<sup>2</sup> | 0.071 ± 0.008 | NA | NA | NA | NA |
| SL12 | 1.50 E<sup>1</sup> | 2.08 E<sup>2</sup> | 1.93 E<sup>2</sup> | 0.072 ± 0.008 | NA | NA | NA | NA |
| SUR | 1.45 E<sup>-2</sup> | 1.84 E<sup>-1</sup> | 1.70 E<sup>-1</sup> | 0.079 ± 0.008 | 3.86 E<sup>-2</sup> | 4.89 E<sup>-1</sup> | 4.50 E<sup>-1</sup> | 0.079 ± 0.003 |
| BW | 1.59 E<sup>0</sup> | 3.34 E<sup>1</sup> | 1.75 E<sup>1</sup> | 0.486 ± 0.036 | 6.69 E<sup>0</sup> | 1.79 E<sup>1</sup> | 1.12 E<sup>1</sup> | 0.373 ± 0.031 |

Note: $\sigma_a^2$ , $\sigma_p^2$ , and $\sigma_e^2$ represent additive genetic variance, phenotype variance, and residual variance; S1-S12, SL1-SL12, SUR and BW: see legend in Table 2; NA, not applicable.

**Figure 2.**
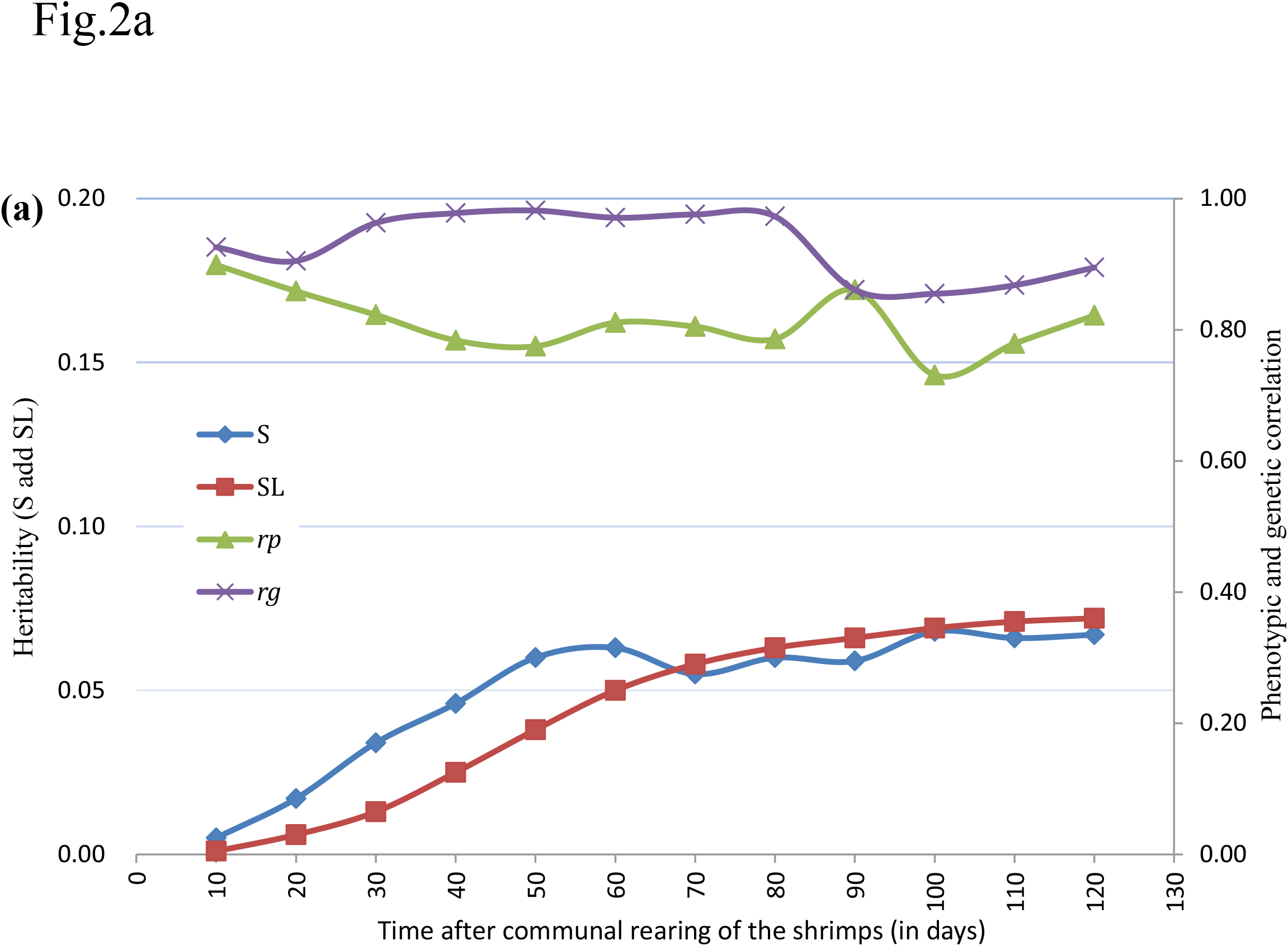
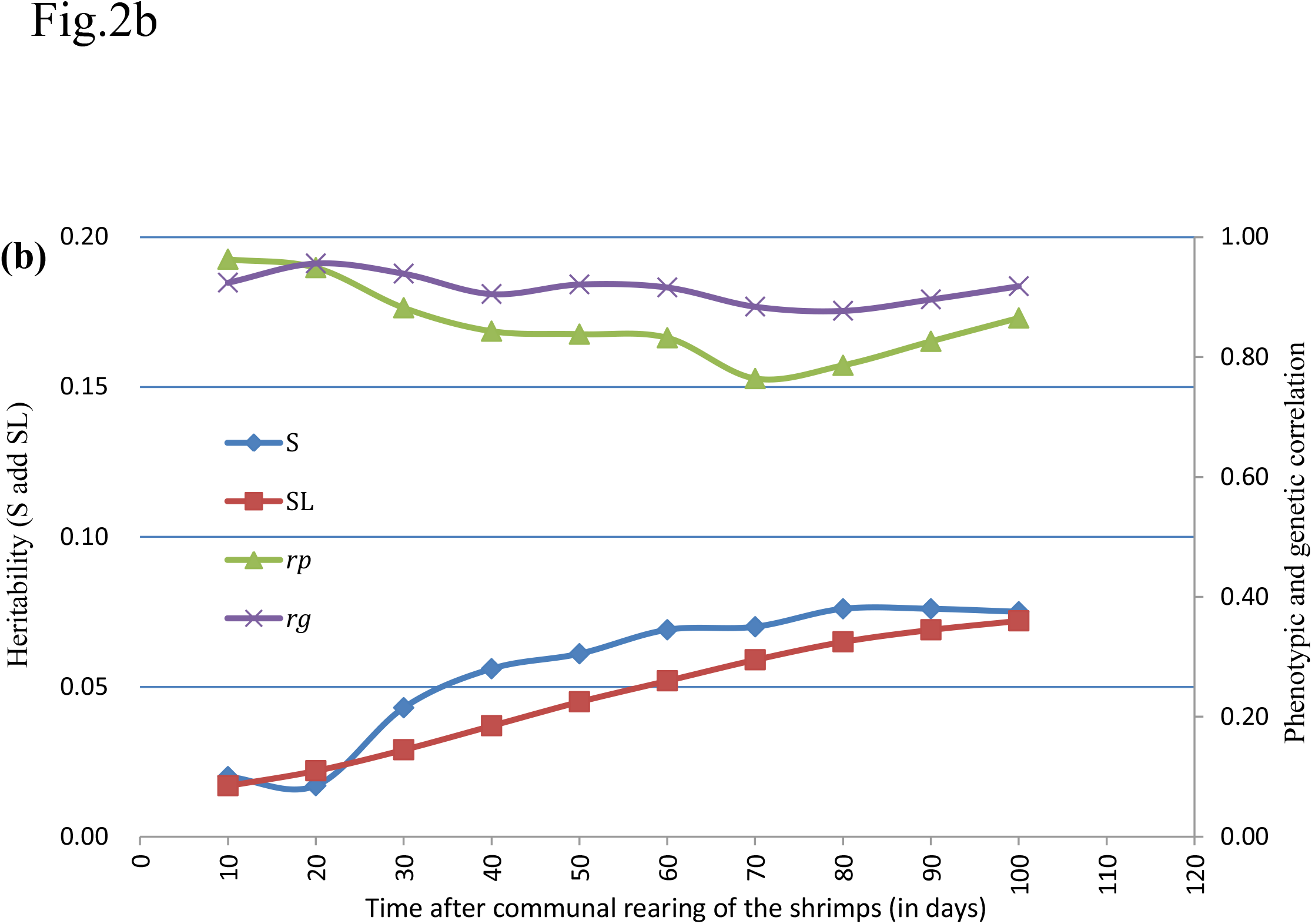
Trajectories of heritability for survival traits (S and SL), phenotypic correlation (*r*_*p*_) and genetic correlation (*r*_*g*_) between S and SL across grow-out test period of (**a**) BL2019 line, (**b**) BL2020 line.

Heritability estimates for final population census survival (SUR) were very similar for S and SL at the final testing stage, while for body weight (BW) in BL2019 and BL2020, they were 0.486 *±* 0.036 and 0.373 *±* 0.031, respectively.

### Phenotype and Genetic Correlations

Summaries of phenotype (*r*_*p*_) and genetic (*r*_*g*_) correlations with SE for S, SL, SUR, and BW are provided in **Table S1-4**. In general *r*_*g*_ for S, estimates between different times were moderate to high and ranged from 0.475 (S1 *vs* S12) to 0.999 (S11 *vs* S12) in the BL2019 breeding line, and 0.277 (S1 *vs* S10) to 0.999 (S9 *vs* S10) for BL2020, respectively. Estimates of *r*_*g*_ for SL between different times were much closer compared with that of S in both two-year breeding lines. As expected, estimates of *r*_*g*_ between SUR and S, or SUR and SL were much lower, and in most cases were only moderate correlations likely due to the issues referred to earlier with unobserved mortality events when estimating SUR vs S/SL. In contrast, estimates of *r*_*g*_ for BW and all survival related traits (S/SL/SUR) were low but all positive. We consider that this result implies that selection directed on survival related traits would not experience potentially negative tradeoff effects on growth traits. Patterns of phenotype correlations (*r*_*p*_) among S, SL, SUR, and BW were similar to those for *r*_*g*_ but were always slightly lower than for *r*_*g*_ (**Figure 3**).

**Figure 3.**
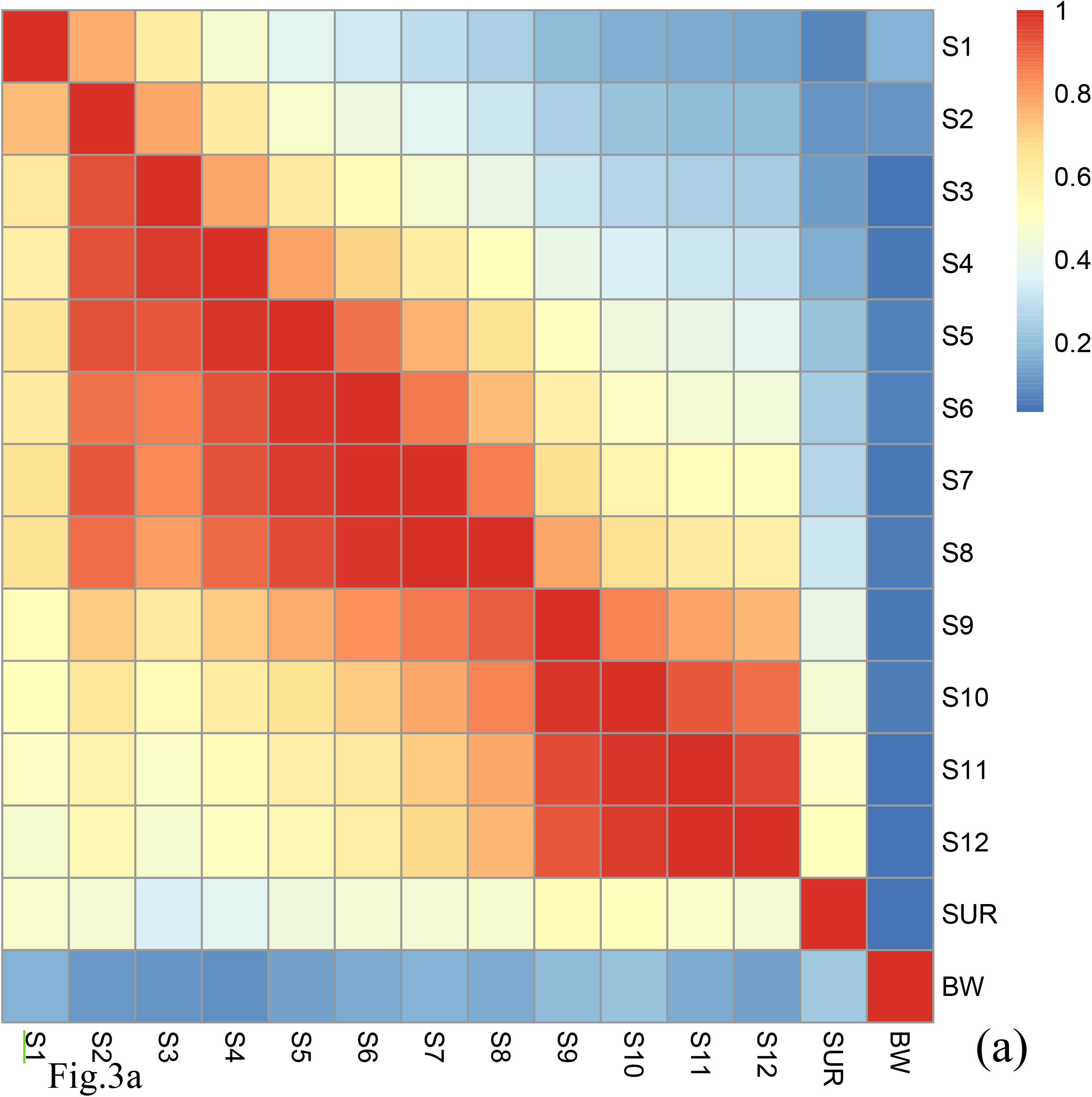
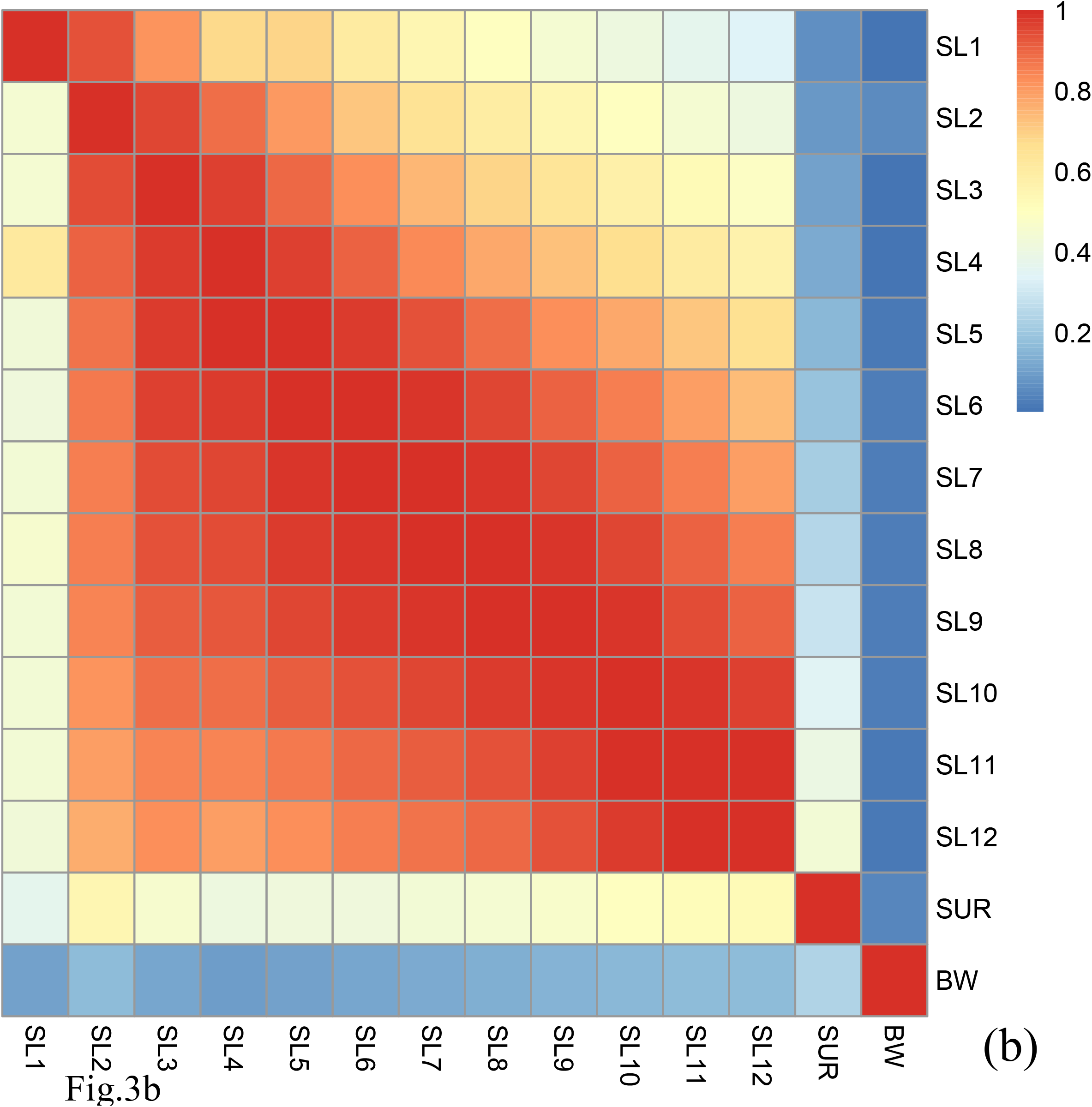
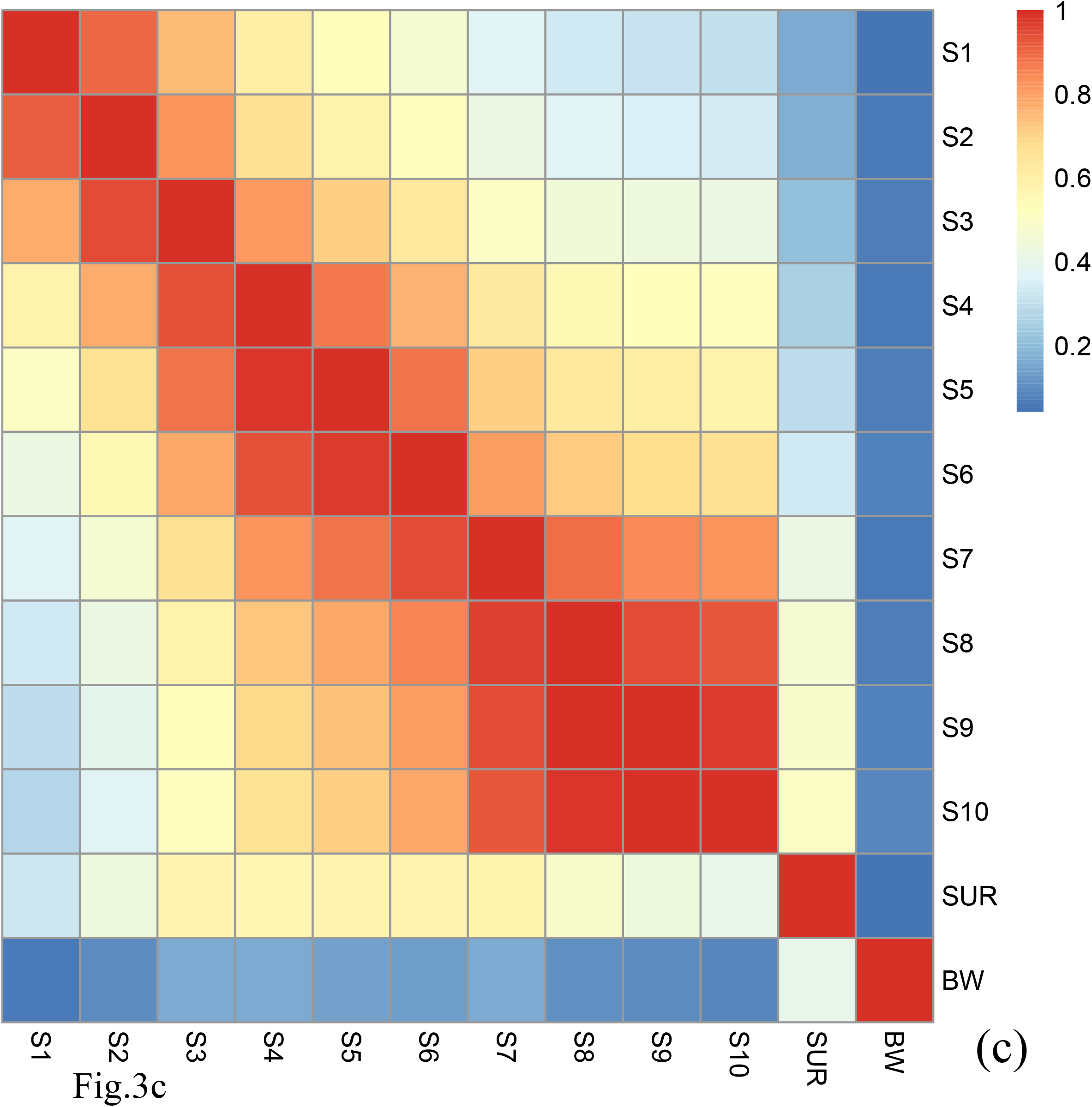
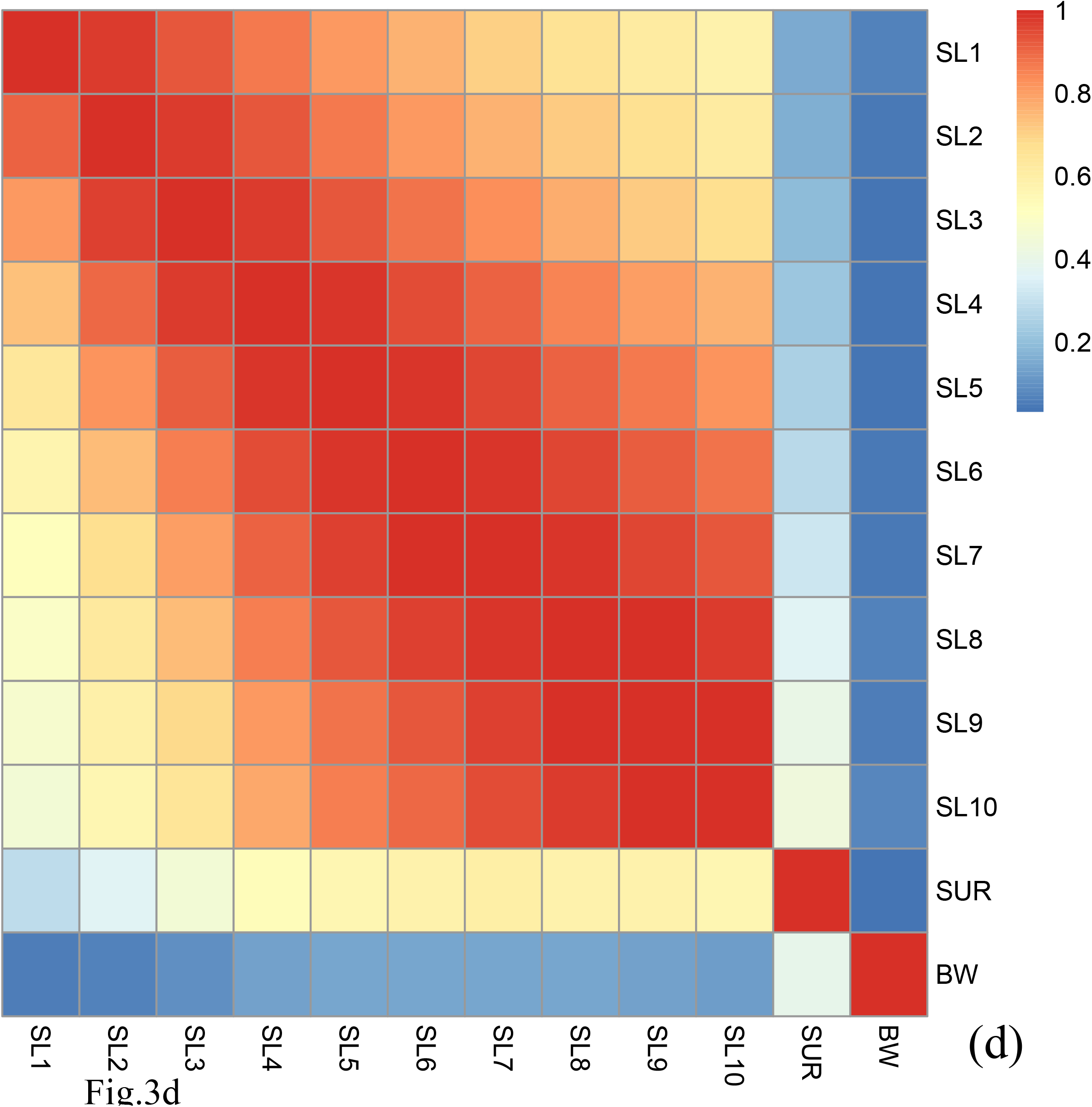
The trajectory of phenotype (above diagonal) and genetic correlations (below diagonal) between **(a)** S1-S12, SUR and BW for the breeding line of BL2019, **(b)** SL1-SL12, SUR and BW of BL2019, **(c)** S1-S10, SUR and BW for the breeding line of BL2020, (**d**) SL1-SL10, SUR and BW for the breeding line of BL2020.

Trajectory of correlations between S and SL estimates across the grow-out test period are illustrated in **Figure 2**. Genetic correlations between binary survival traits (S) and survival days (SL) were all high at the different observation times and ranged from 0.855 to 0.982 for BL2019 and 0.877 to 0.956 for BL2020. The results reported above suggest that families with high survival rates are also likely to show a general trend for a longer mean life span over the test period. Trajectories of genetic correlations for S traits across time were consistent with that of estimation for SL, with estimates for *r*_*g*_ gradually decreasing across the experimental time period (**Figure 3**). Patterns of change on *r*_*g*_ estimates for S/SL across time were similar between breeding lines in both years.

### Full-sib Family Effects

The likelihood ratio test suggested that there were very limited full-sib family effects (*c*^*2*^) in the genetic analysis. Among 26 comparison for BL2019, 24 tests were not significant for the likelihood ratio tests (**Table 5S**). In contrast, S1 and SUR were significant for *c*^*2*^ (*χ*^*2*^_*1df*_). Data interpretation for genetic parameters for S1 and SUR however, showed limited potential for improving an animal model fit for full-sib family effects (**Model 3**) because the new results for S1 and SUR genetic parameters showed large associated SE estimates (S1, *h*^*2*^ = 0.005 ± 0.009; SUR, *h*^*2*^ = 0.053 ± 0.070). Similarly, there was limited potential for improving animal model fit for full-sib family effects in the genetic analysis of survival data from BL2020 based on the result of likelihood ratio tests.

## 4 DISCUSSION

The current study provides the first genetic analysis of the trajectory in survival traits across time for improved lines in a farmed aquatic species. Results of the genetic evaluation indicate that heritability estimates for the two defined types survival applied here were very similar. In general, estimated levels of heritability for survival traits were low but increased gradually across the grow-out test periods for both year classes. In addition, genetic correlations between the binary trait of survivability (S) and the continuous trait of survival time (SL) were relatively high. These findings will be useful for genetic analyses of survival data in field environments and potentially contribute to the development of relatively robust farmed aquatic strains used in aquaculture. An issue we identified in the analysis however, was missing records of some individual mortality events across our test periods. This reflects in general the difficulty of collecting survival data for aquaculture species in the field when working in commercial test environments.

### Mortality

Records of daily mortality in breeding lines are not only important for genetic evaluation of survival traits, but also for assessment of general husbandry management of population health status in breeding lines. Changes in daily mortality here were similar to those reported in our previous study with the mortality rate varying between 0.1% - 0.15% (Ren et al., 2020a). Designing and managing reliable, high quality water culture conditions is a major constraint in selective breeding programs for penaeid shrimps. Most daily mortality rates for farmed penaeids domestication programs range between 0.1% - 0.5%, depending on the different types of culture systems employed (Coman et al., 2005; Duy et al., 2012; Yano, 2000).

The highest daily mortality peaks in both test years occurred in late grow-out stages (**Figure 1**), periods that coincided with the coldest weather temperatures in the winter seasons at the hatchery location in China. Physical water temperature parameters ranged from 22 to 29 °C which provides ideal conditions to reach pre-maturation stage in white-leg shrimp. In contrast, temperature records for the peaks in daily morality during winter ranged from 17-19 °C. In general, husbandry management of penaeid breeding lines are considered appropriate if daily mortality rate fluctuates around 0.5%, and higher mortality rates do not last for longer than 7-10 days. In our study when mean daily temperatures increased from this relatively low range, mortality decreased to normal levels again. Furthermore, in most cases individuals that died during low temperature periods were of small size and were weak individuals.

In the current study, we observed a gap in bias between survival estimators across the test period and the final population census data. Collecting survival data in natural ‘field’ conditions can be difficult in aquaculture breeding programs compared with that in terrestrial farm animals because most mortality events are under water and consequently, may not be detected. While survival phenotype derived from controlled experimental challenge tests can be more accurate (Ødegård et al., 2011 ; Yáñez et al., 2014), the use of survival data in natural field conditions can often provide better information for developing robust culture lines in aquaculture species (Bangera et al., 2014; Dégremont et al., 2015; Houston et al., 2008; Lillehammer et al., 2013). In particular, survival data collected under field conditions will better reflect natural processes of changes in mortality in a population and collecting these data can avoid potential G-by-E impacts when using artificial challenge test environments.

Survival estimators and final population census results in the current study were comparable with earlier reports on genetic breeding programs conducted on penaeid shrimps. In Mexico, survival rate of *P. vannamei* selection lines were reported to range from 71% to 82.2% during the grow-out test stages (Campos-Montes et al., 2013; Caballero-Zamora et al., 2015). Breeding nucleus data for farmed *P. vannamei* in Colombia reported survival rates ranging from 56.9% to 77.2% (Gitterle et al., 2005) while they were 70% and 73% for two grow-out stages in a selection program for *P. monodon* in Australia (Coman et al., 2010). In Vietnam, survival rates for *P. monodon* were 34% to 49% in a family selection program (Van et al., 2020).

### Animal Models for Survival Analysis

While survival traits are often recorded as non-normally distributed binary traits, reports of genetic evaluation of survival data in dairy cattle (Sasaki et al., 2015; Van Pelt et al., 2015, 2016; Heise et al., 2016, 2018) suggest that linear models are likely to be more appropriate for genetic analysis of survival data in aquaculture. Proportion hazards models (PHM) have been often considered to fit time-to-event survival data better (Ducrocq, 1994; Neerhof et al., 2000), however, computing time to analyse these data successfully is much high than with linear models. Furthermore, using a PHM approach it is more difficult to assess genetic correlations with other continuous traits. Similarly, threshold models require significantly more computing resources for genetic analysis of survival data compared with linear models. Moreover, correlations of EBVs estimated with threshold models and linear models have been almost identical, indicating that there are few advantages of applying threshold models in routine genetic analyses of survival data (Boettcher et al., 1999; Vazquez et al., 2009). In practical terms, consideration of model choice in breeding programs can be a balance between considerations of multiple factors, including: simplified data records, comparative model performance, available computing resources, and integration of data from other important economic traits at the same time. We therefore consider linear multivariate animal models (LMA) to provide informative and reliable tools for routine genetic analysis of survival data in farmed aquatic species.

### Heritability Estimates

Data structure in the current study applied a larger number of full paternity and half-sib families that were suitable for achieving highly accurate heritability estimates. In addition, full and half-sib families were produced over a relatively short time period (one week) that effectively removes time of age effects and common environmental effects in the genetic analysis. This can enhance the accuracy of the animal model results generated. We therefore chose a linear multivariate animal model (LMA) approach for our genetic analyses rather than other more sophisticated linear animal models such as a random regression model or repeatability model to fit age effects of time for the survival data set here.

Heritability estimates for S1 to S12 and SL1 to SL 12 were consistent with results reported from other white-leg shrimp breeding programs. Gitterle et al. (2005) evaluated binary survival traits based on 430 full-sib families in different test environments and reported heritability estimates that ranged from 0.04 to 0.10. In another genetic analysis of 2008 to 2010 survival data of white-leg shrimp in Mexico, heritability estimates for early stage survival were 0.03 and 0.04 for later grow-out test stage (Campos-Montes et al., 2013). In a G-by-E effect study, heritability of survival was reported to be 0.06 under cold temperature conditions, while it was much higher (0.11) at normal ambient temperatures (Li et al., 2015). In addition, genetic parameters for survival in the presence or absence of a white spot disease outbreak were reported to be 0.00 and 0.06 (Caballero-Zamora et al., 2015), respectively. Overall, these results for heritability estimates of survival traits agree well with quantitative genetic theoretical predictions that survival traits are a group of fitness-associated traits that tend to show the lowest levels of heritability (Hill, 2010).

Here we compared two types of survival definitions; binary traits and continuous traits for our genetic evaluation, with both showing similar heritability estimates (**Table 3**). While this result did not fit our earlier assumption that continuous traits would improve the results for heritability estimates via adding more survival time information to mortality events, similar findings have been reported for genetic analysis of survival data in experimental challenge tests on aquaculture species (Joshi et al., 2021a; Vu et al., 2021). Binary trait recording however, is much more simple (0, 1) to score than continuous survival data in practice and thus, should be considered more feasible for routine genetic evaluation in aquaculture breeding programs.

Knowledge about heritability estimate trajectories across time is important to improve efficiency in selection programs and to provide insights on predicted selection response over the time period in the test. We believe that our study is the first report on the heritability of survival trajectory across time in a farmed aquatic species. Results here, suggest that for both lines and applying both types of survival trait definition, similar patterns for trajectory with time were evident. In general, initial heritability estimates were almost close to 0, following which they gradually increased across the grow-out test period before slowing down or reaching a plateau during the latter stages (**Figure 2**).

Of interest, this pattern in trajectory is similar compared with some earlier reports for changes in growth heritability estimates across different age times (Turra et al., 2012; He et al., 2018). In dairy science, similar patterns for survival trait heritability were also reported for a 72 month period of milk production life (Van Pelt et al., 2015).

Common full-sib effects (*c*^*2*^) were not significant in the current study indicating that applying a linear multivariate animal model (LMA) without full-sib effects was effective for routine genetic evaluation here. This finding agrees with a systematic review paper on *c*^*2*^ estimation in aquaculture breeding that suggests that full-sib effects contribute only a small proportion of total phenotypic variance at earlier growth stages for growth related traits, but for growth traits in an individual’s later stages or with other phenotypic traits, full-sib effects (*c*^*2*^) are not significant in most cases and are essentially zero (Nguyen, 2021). As an example routine genetic evaluation models applied in commercial tilapia selection breeding programs based on current published genetic animal models do not include full-sib effects (*c*^*2*^) (Joshi et al., 2021 b).

### Genetic and Phenotype Correlations

Both genetic and phenotypic correlations between binary and continuous survival traits time showed high correlations (>0.8) at each time window (**Figure 2**). In an aquaculture context, similar patterns have been reported for correlations between different types of growth related traits, including for *r*_*g*_*/r*_*p*_ among body weight, body length, and other morphological growth traits (He et al., 2018; Schlicht et al., 2018). Patterns for *r*_*g*_*/r*_*p*_ also support findings from heritability estimates that both survival definitions can be applied equally well for genetic selection on overall survival trait data in commercial breeding lines.

The current study is the first report on patterns of genetic correlations for survival traits across times in a farmed aquatic species (**Figure 3**). In general, patterns of genetic correlation for survival traits across time (*r*_*g*_ among S1 to S12 and SL1 to SL12) in the current study were moderate to high. There was a trend here for a gradual decrease in *r*_*g*_ between survival traits with the time. These patterns for *r*_*g*_ were in accordance with trajectories of genetic correlations for growth traits across ages in other aquaculture studies (Turra et al., 2012; He et al., 2018; Schlicht et al., 2018), as well as for survival traits in livestock (Van Pelt et al., 2015). In addition, results for *r*_*p*_ trajectories of correlation across time between S1 to S12 and/or SL1 to SL12 were similar with that of *r*_*g*_, but values for *r*_*p*_ are always slightly lower than for *r*_*g*_ (**Figure 3**).

In contrast, genetic correlations between SUR and S1-12/SL1-12 were much lower but nevertheless, in most instances were still moderate (**Figure 3**). From observation of the patterns of survival traits across times, the real genetic correlations between SUR and other survival traits are likely to be much higher than that observed here. This most likely resulted from the missing data of survival records during the grow-out period and the final population census. Highly accurate recording of survival data in field commercial aquaculture environments however, is often difficult compared with data collection in challenge test experimental conditions, particularly, for white-leg shrimp that are of relatively small size and that are cannibalistic. These characteristics mean that some results of mortality events during grow-out test period are unlikely ever to be detected under field conditions.

In the current study, correlations between body weight and survival traits were generally low (**Figure 3**), but still positive, a result that is consistent with previous work on white-leg shrimp (Campos-Montes et al., 2013; Caballero-Zamora et al., 2015; Li et al., 2015; Ren et al., 2020a). This suggests overall, that survival and growth are two groups of independent traits and can be selected for simultaneously in a genetic breeding program via a multi-traits selection approach.

### Implications

While survival can be considered a relatively ‘old’ trait in animal breeding, often neglected because of the simple records (0 or 1) scored and generally low heritability estimates. Developing routine genetic evaluation systems for survival traits however, will be important for developing robust strains using valuable field survival data. Firstly, survival data collection from commercial lines can be used directly for improving overall survival rate in target breeding lines (Gjedrem and Rye, 2018). Moreover, survival data recorded from commercial lines when special events arise e.g. disease outbreaks can also be applied effectively to developing disease resistant strains, and these type of data are also valuable for avoiding potential G-by-E effects compared with survival data from controlled (artificial) challenge experiments (Bangera et al., 2014; Dégremont et al., 2015; Barría et al., 2020; Fraslin et al., 2022). Additionally, routine genetic evaluation of survival data in breeding lines provides a vital assessment of the relative health status of a stock under the general husbandry management practices employed on the breeding lines. Here we provide the first report of the trajectory of heritability estimates for survival traits across the grow-out period for white-leg shrimp breeding lines in China. The results will be useful for applying commercial field survival data to developing improved robust culture lines of this important crustacean species in the future.

## 5 CONCLUSIONS

In conclusion, we developed routine genetic evaluations for survival data in improved white-leg shrimp culture lines in China. Results of heritability and genetic correlation estimates indicate that both survival definitions for binary traits and continuous traits of survival time can be used effectively for genetic analysis of survival data. Binary survival records following with linear multivariate animal models (LMA) provide a feasible routine genetic evaluation approach in practical commercial breeding programs because of the simple data recording, reduced computing time required, and the ability to combine performance assessment of multiple continuous phenotypic traits in the genetic analysis. While heritability estimates for survival traits here were generally low, they showed a gradual increasing trend across the grow-out period. Genetic correlations for survival traits during grow-out tests were moderate to high, and the closer the times were between estimates of two survival traits, the higher were the genetic correlations. Genetic correlations between survival traits and body weight were low but all estimates were positive. In summary, we first reported the trajectory of heritability estimates for survival traits across time in aquaculture genetics, which would be useful for how to use survival data in field situations to develop more robust culture strains of *P. vannamei* and potentially for other farmed aquatic species in the future.

## 7 DATA AVAULABILITY STATEMENT

The raw data used to support the findings of the current research are available from the corresponding author upon request.

## 8 AUTHOR CONTRIBUTIONS

S.J.R. and D.A.H. conceptualized the research, developed animal models for data analyses. S.J.R. and P.B.M. wrote the manuscript. All authors have critically revised the manuscript.

## 10 ACKNOWLEDGEMENT

The authors would like to thank Ms. Haixiang Lan for collecting the survival data used in this study and thank Mr. Zhante Shi for technical support during the grow-out testing of the shrimp breeding lines.

## Additional files

**Additional file 1 Table S1**

Format: csv file

Title: Estimated genetic (below diagonal) and phenotypic correlations (above diagonal) between S1-S12, SUR and BW for the BL2019 breeding line (estimates ± se).

**Additional file 2 Table S2**

Format: csv file

Title: Estimated genetic (below diagonal) and phenotypic correlations (above diagonal) between SL1-SL12, SUR and BW for the BL2019 breeding line (estimates ± se).

**Additional file 3 Table S3**

Format: csv file

Title: Estimated genetic (below diagonal) and phenotypic correlations (above diagonal) between S1-S10, SUR and BW for the BL2020 breeding line (estimates ± se).

**Additional file 4 Table S4**

Format: csv file

Title: Estimated genetic (below diagonal) and phenotypic correlations (above diagonal) between SL1-SL10, SUR and BW for the BL2020 breeding line (estimates ± se).

**Additional file 5 Table S5**

Format: docx file

Table S5. Univariate genetic analyses of the likelihood ratios and the full-sib family effect (*c*^*2*^± se) for BL2019 breeding line, significance test at *α* = 0.05 (*χ*^*2*^ _1DF_).

## REFFERENCES

Bangera, R., Ødegård, J., Mikkelsen, H., Nielsen, H. M., Seppola, M., Puvanendran, V., et al. (2014). Genetic analysis of francisellosis field outbreak in Atlantic cod (Gadus morhua L.) using an ordinal threshold model. Aquaculture 420, S50–S56.

Barría, A., Christensen, K. A., Yoshida, G., Jedlicki, A., Leong, J. S., Rondeau, E. B., et al. (2019). Whole genome linkage disequilibrium and effective population size in a coho salmon (Oncorhynchus kisutch) breeding population using a high-density SNP array. Front. Genet. 10, 498.

Barría, A., Trinh, T. Q., Mahmuddin, M., Benzie, J. A., Chadag, V. M., and Houston, R. D. (2020). Genetic parameters for resistance to Tilapia Lake Virus (TiLV) in Nile tilapia (Oreochromis niloticus). Aquaculture 522, 735126.

Bernatchez, L., Wellenreuther, M., Araneda, C., Ashton, D. T., Barth, J. M., Beacham, T. D., et al. (2017). Harnessing the power of genomics to secure the future of seafood. Trends Ecol. Evol. 32(9), 665–680.

Boettcher, P. J., Jairath, L. K., and Dekkers, J. C. M. (1999). Comparison of methods for genetic evaluation of sires for survival of their daughters in the first three lactations. J. Dairy Sci. 82(5), 1034–1044.

Caballero-Zamora, A., Montaldo, H. H., Campos-Montes, G. R., Cienfuegos-Rivas, E. G., Martínez - Ortega, A., et al. (2015). Genetic parameters for body weight and survival in the Pacific White Shrimp Penaeus (Litopenaeus) vannamei affected by a White Spot Syndrome Virus (WSSV) natural outbreak. Aquaculture 447, 102–107.

Campos-Montes, G. R., Montaldo, H. H., Martínez -Ortega, A., Jiménez, A. M., et al. (2013). Genetic parameters for growth and survival traits in Pacific white shrimp Penaeus (Litopenaeus) vannamei from a nucleus population undergoing a two-stage selection program. Aquac. Int. 21(2), 299–310.

Coman, G. I., Crocos, P. I., Arnold, S. J., Keys, S. I., Preston, N. P., et al. (2005). Growth, survival and reproductive performance of domesticated Australian stocks of the giant tiger prawn, Penaeus monodon, reared in tanks and raceways. J. World Aquac. Soc. 36(4), 464–479.

Coman, G. J., Arnold, S. J., Wood, A. T., and Kube, P. D. (2010). Age: age genetic correlations for weight of Penaeus monodon reared in broodstock tank systems. Aquaculture 307(1-2), 1–5.

Correa, K., Lhorente, J. P., López, M. E., Bassini, L., Naswa, S., Deeb, N., et al. (2015). Genome-wide association analysis reveals loci associated with resistance against Piscirickettsia salmonis in two Atlantic salmon (Salmo salar L.) chromosomes. BMC Genom. 16(1), 1–9.

Dégremont, L., Garcia, C., and Allen Jr, S. K. (2015). Genetic improvement for disease resistance in oysters: a review. J. Invertebr. Pathol. 131, 226–241.

Ducrocq, V. (1994). Statistical analysis of length of productive life for dairy cows of the Normande breed. J. Dairy Sci. 77(3), 855–866.

Ducrocq, V., and Sölkner, J. (1994). “The survival kit”: a fortran package for the analysis of survival data. Pages 51–52 in Proc.5th World Congr. Genet. Appl. Livest. Prod., Guelph, Ontario, Canada.

Ducrocq, V., and Casella, G. (1996). A Bayesian analysis of mixed survival models. Genet. Sel. Evol. 28(6), 505–529.

Ducrocq, V., Sölkner, J., and Mészáros, G. (2010). Survival Kit V6 —A software package for survival analysis. 9th World Congr. Genet. Appl. Livest. Prod., Leipzig, Germany.

Duy, H. N., Coman, G. J., Wille, M., Wouters, R., Quoc, H. N., Vu, T., et al. (2012). Effect of water exchange, salinity regime, stocking density and diets on growth and survival of domesticated black tiger shrimp Penaeus monodon (Fabricius, 1798) reared in sand-based recirculating systems. Aquaculture 338, 253–259.

FAO, 2020. Aquaculture Production (Quanlities and values). FishStatJ -Software for Fishery Statistical Time Series. FAO Fisheries and Aquaculture Department, Rome.

Forabosco, F., Jakobsen, J. H., and Fikse, W. F. (2009). International genetic evaluation for direct longevity in dairy bulls. J. Dairy Sci. 92(5), 2338–2347.

Fraslin, C., Koskinen, H., Nousianen, A., Houston, R. D., and Kause, A. (2022). Genome-wide association and genomic prediction of resistance to Flavobacterium columnare in a farmed rainbow trout population. Aquaculture 738332.

Friggens, N. C., Blanc, F., Berry, D. P., and Puillet, L. (2017). Deciphering animal robustness. A synthesis to facilitate its use in livestock breeding and management. Animal 11(12), 2237–2251.

Gitterle, T., Rye, M., Salte, R., Cock, J., Johansen, H., Lozano, C., et al. (2005). Genetic (co) variation in harvest body weight and survival in Penaeus (Litopenaeus) vannamei under standard commercial conditions. Aquaculture 243(1-4), 83–92.

Gjedrem, T., Robinson, N., and Rye, M. (2012). The importance of selective breeding in aquaculture to meet future demands for animal protein: a review. Aquaculture 350, 117–129.

Gjedrem, T., and Rye, M. (2018). Selection response in fish and shellfish: a review. Rev. Aquac. 10(1), 168–179.

Gutierrez, A. P., Bean, T. P., Hooper, C., Stenton, C. A., Sanders, M. B., Paley, R. K., et al. (2018). A genome-wide association study for host resistance to ostreid herpesvirus in Pacific oysters (Crassostrea gigas). G3: Genes, Genom. Genet. 8(4), 1273–1280.

He, J., Zhao, Y., Zhao, J., Gao, J., Han, D., Xu, P., and Yang, R. (2017). Multivariate random regression analysis for body weight and main morphological traits in genetically improved farmed tilapia (Oreochromis niloticus). Genet. Sel. Evol. 49(1), 1–13.

Heise, J., Liu, Z., Stock, K. F., Rensing, S., Reinhardt, F., and Simianer, H. (2016). The genetic structure of longevity in dairy cows. J. Dairy Sci. 99(2), 1253–1265.

Heise, J., Stock, K. F., Reinhardt, F., Ha, N. T., and Simianer, H. (2018). Phenotypic and genetic relationships between age at first calving, its component traits, and survival of heifers up to second calving. J. Dairy Sci. 101(1), 425–432.

Hung, D., Nguyen, N. H., Hurwood, D. A., and Mather, P. B. (2013). Quantitative genetic parameters for body traits at different ages in a cultured stock of giant freshwater prawn (Macrobrachium rosenbergii) selected for fast growth. Mar. Freshw. Res. 65(3), 198–205.

Hill, W. G. (2010). Understanding and using quantitative genetic variation. Philos. Trans. R. Soc. B: Biol. Sci. 365(1537), 73–85.

Houston, R. D. (2017). Future directions in breeding for disease resistance in aquaculture species. Rev. Bras. de Zootec. 46, 545–551.

Houston, R. D., Bean, T. P., Macqueen, D. J., Gundappa, M. K., Jin, Y. H., Jenkins, T. L., et al. (2020). Harnessing genomics to fast-track genetic improvement in aquaculture. Nat. Rev. Genet. 21(7), 389–409.

Houston, R. D., Haley, C. S., Hamilton, A., Guy, D. R., Tinch, A. E., Taggart, J. B., et al. (2008). Major quantitative trait loci affect resistance to infectious pancreatic necrosis in Atlantic salmon (Salmo salar). Genetics 178(2), 1109–1115.

Joshi, R., Skaaurd, A., and Tola Alvarez, A. (2021a). Experimental validation of genetic selection for resistance against Streptococcus agalactiae via different routes of infection in the commercial Nile tilapia breeding programme. J. Anim. Breed. Genet. 138(3), 338–348.

Joshi, R., Skaarud, A., Alvarez, A. T., Moen, T., and Ødegård, J. (2021b). Bayesian genomic models boost prediction accuracy for survival to Streptococcus agalactiae infection in Nile tilapia (Oreochromus nilioticus). Genet. Sel. Evol. 53(1), 1–10.

Kaplan, E. L., and Meier, P. (1958). Nonparametric estimation from incomplete observations. J. Am. Stat. Assoc. 53(282), 457–481.

Kassambara, A., Kosinski, M., Biecek, P., and Fabian, S. (2017). Package ‘survminer’. Drawing Survival Curves using ‘ggplot2’. R package version 0.3.1.

Kumar, G., and Engle, C. R. (2016). Technological advances that led to growth of shrimp, salmon, and tilapia farming. Rev. Fish. Sci. Aquac. 24(2), 136–152.

Li, W., Luan, S., Luo, K., Sui, J., Xu, X., Tan, J., et al. (2015). Genetic parameters and genotype by environment interaction for cold tolerance, body weight and survival of the Pacific white shrimp Penaeus vannamei at different temperatures. Aquaculture 441, 8–15.

Lillehammer, M., Ødegård, J., Madsen, P., Gjerde, B., Refstie, T., and Rye, M. (2013). Survival, growth and sexual maturation in Atlantic salmon exposed to infectious pancreatic necrosis: a multi-variate mixture model approach. Genet. Sel. Evol. 45(1), 1–12.

Meyer, K. (2007). WOMBAT—A tool for mixed model analyses in quantitative genetics by restricted maximum likelihood (REML). J. Zhejiang Univ. Sci. B 8(11), 815–821.

Naylor, R. L., Hardy, R. W., Buschmann, A. H., Bush, S. R., Cao, L., Klinger, D. H., et al. (2021). A 20-year retrospective review of global aquaculture. Nature 591(7851), 551–563.

Neerhof, H. J., Madsen, P., Ducrocq, V. P., Vollema, A. R., Jensen, J., and Korsgaard, I. R. (2000). Relationships between mastitis and functional longevity in Danish Black and White dairy cattle estimated using survival analysis. J. Dairy Sci. 83(5), 1064–1071.

Nguyen, N. H. (2016). Genetic improvement for important farmed aquaculture species with a reference to carp, tilapia and prawns in Asia: achievements, lessons and challenges. Fish Fish. 17(2), 483–506.

Nguyen, N. H. (2021). A systematic review and meta-analysis of genetic parameters for complex quantitative traits in aquatic animal species. bioRxiv doi.org/10.1101/2021.05.20.445048

Ødegård, J., Olesen, I., Gjerde, B., and Klemetsdal, G. (2006). Evaluation of statistical models for genetic analysis of challenge test data on furunculosis resistance in Atlantic salmon (Salmo salar): Prediction of field survival. Aquaculture 259(1-4), 116–123.

Ødegård, J., Baranski, M., Gjerde, B., and Gjedrem, T. (2011). Methodology for genetic evaluation of disease resistance in aquaculture species: challenges and future prospects. Aquac. Res. 42, 103–114.

Palaiokostas, C., Cariou, S., Bestin, A., Bruant, J. S., Haffray, P., Morin, T., et al. (2018). Genome-wide association and genomic prediction of resistance to viral nervous necrosis in European sea bass (Dicentrarchus labrax) using RAD sequencing. Genet. Sel. Evol. 50(1), 1–11.

R Core Team (2019). R: A language and environment for statistical computing. Vienna, Austria.

Reid, G. K., Gurney-Smith, H. J., Marcogliese, D. J., Knowler, D., Benfey, T., Garber, A. F., et al. (2019). Climate change and aquaculture: considering biological response and resources. Aquac. Environ. Interact. 11, 569–602.

Ren, S., Mather, P. B., Tang, B., and Hurwood, D. A. (2018). Levels of genetic diversity and inferred origins of Penaeus vannamei culture resources in China: Implications for the production of a broad synthetic base population for genetic improvement. Aquaculture 491, 221–231.

Ren, S., Prentis, P., Mather, P. B., Li, Y., Tang, B., and Hurwood, D. A. (2020a). Genetic parameters for growth and survival traits in a base population of Pacific white shrimp (Litopenaeus vannamei) developed from domesticated strains in China. Aquaculture 523, 735148.

Ren, S., Mather, P. B., Prentis, P., Li, Y., Tang, B., and Hurwood, D. A. (2020b). Quantitative genetic assessment of female reproductive traits in a domesticated pacific white shrimp (Penaeus vannamei) line in China. Sci. Rep. 10(1), 1–10.

Ren, S., Mather, P. B., Tang, B., and Hurwood, D. A. (2020c). Comparison of reproductive performance of domesticated Litopenaeus vannamei females reared in recirculating tanks and earthen ponds: an evaluation of reproductive quality of spawns in relation to female body size and spawning order. Front. Mar. Sci. 7, 560.

Ren, S., Mather, P. B., Tang, B., and Hurwood, D. A. (2022). Standardized microsatellite panels for pedigree management of farmed white-leg shrimp (Penaeus vannamei) stocks validated in a VIE tagged family selection line. Aquaculture 551, 737946.

Sasaki, O., Aihara, M., Nishiura, A., Takeda, H., and Satoh, M. (2015). Genetic analysis of the cumulative pseudo-survival rate during lactation of Holstein cattle in Japan by using random regression models. J. Dairy Sci. 98(8), 5781–5795.

Schaeffer, L. R. (2004). Application of random regression models in animal breeding. Livest. Prod. Sci. 86(1-3), 35–45.

Schlicht, K., Krattenmacher, N., Lugert, V., Schulz, C., Thaller, G., and Tetens, J. (2018). Genetic analysis of production traits in turbot (Scophthalmus maximus) using random regression models based on molecular relatedness. J. Anim. Breed. Genet. 135(4), 275–285.

Sewalem, A., Miglior, F., and Kistemaker, G. J. (2010). Analysis of the relationship between workability traits and functional longevity in Canadian dairy breeds. J. Dairy Sci. 93(9), 4359–4365.

Suebsong, W., Poompuang, S., Srisapoome, P., Koonawootrittriron, S., Luengnaruemitchai, A., Johansen, H., et al. (2019). Selection response for Streptococcus agalactiae resistance in Nile tilapia Oreochromis niloticus. J. Fish Dis. 42(11), 1553–1562.

Tarrés, J., Bidanel, J. P., Hofer, A., and Ducrocq, V. (2006). Analysis of longevity and exterior traits on Large White sows in Switzerland. J. Anim Sci. 84(11), 2914–2924.

Trang, T. T., Hung, N. H., Ninh, N. H., Knibb, W., and Nguyen, N. H. (2019). Genetic variation in disease resistance against White Spot Syndrome Virus (WSSV) in Liptopenaeus vannamei. Front. Genet. 10, 264.

Troell, M., Naylor, R. L., Metian, M., Beveridge, M., Tyedmers, P. H., Folke, C., et al. (2014). Does aquaculture add resilience to the global food system? Proc. Nati. Acad. Sci. 111(37), 13257–13263.

Turra, E. M., de Oliveira, D. A. A., Valente, B. D., de Alencar Teixeira, E., de Assis Prado, S., de Melo, D. C., et al. (2012). Estimation of genetic parameters for body weights of Nile tilapia Oreochromis niloticus using random regression models. Aquaculture 354, 31–37.

Vallejo, R. L., Leeds, T. D., Gao, G., Parsons, J. E., Martin, K. E., Evenhuis, J. P., et al. (2017). Genomic selection models double the accuracy of predicted breeding values for bacterial cold water disease resistance compared to a traditional pedigree-based model in rainbow trout aquaculture. Genet. Sel. Evol. 49(1), 1–13.

Van Pelt, M. L., Meuwissen, T. H. E., De Jong, G., and Veerkamp, R. F. (2015). Genetic analysis of longevity in Dutch dairy cattle using random regression. J. Dairy Sci. 98(6), 4117–4130.

Van Pelt, M. L., Ducrocq, V., De Jong, G., Calus, M. P. L., and Veerkamp, R. F. (2016). Genetic changes of survival traits over the past 25 yr in Dutch dairy cattle. J. Dairy Sci. 99(12), 9810–9819.

Van Sang, N., Luan, N. T., Van Hao, N., Van Nhien, T., Vu, N. T., and Nguyen, N. H. (2020). Genotype by environment interaction for survival and harvest body weight between recirculating tank system and pond culture in Penaeus monodon. Aquaculture 525, 735278.

Vazquez, A. I., Gianola, D., Bates, D., Weigel, K. A., and Heringstad, B. (2009). Assessment of Poisson, logit, and linear models for genetic analysis of clinical mastitis in Norwegian Red cows. J Dairy Sci. 92(2), 739–748.

Veerkamp, R. F., Engel, B., and Brotherstone, S. (2002). Single and multitrait estimates of breeding values for survival using sire and animal models. Anim. Sci. 75(1), 15–24.

Vu, N. T., Phuc, T. H., Oanh, K. T. P., Sang, N. V., Trang, T. T., and Nguyen, N. H. (2022). Accuracies of genomic predictions for disease resistance of striped catfish to Edwardsiella ictaluri using artificial intelligence algorithms. G3: Genes, Genom. Genet. 12(1), jkab361.

Yáñez, J. M., Houston, R. D., and Newman, S. (2014). Genetics and genomics of disease resistance in salmonid species. Front. Genet. 5, 415.

Yano, I. (2000). Cultivation of broodstock in closed recirculating system in specific pathogen free (SPF) penaeid shrimp. Aquac. Sci. 48(2), 249–257.

Zavadilová, L., Němcová, E., and Štípková, M. (2011). Effect of type traits on functional longevity of Czech Holstein cows estimated from a Cox proportional hazards model. J. Dairy Sci. 94(8), 4090–4099.

