## Supplemental Table 5 for "Insight on Selective Breeding the Robustness Based on Field Survival Records: New Genetic Evaluation of Survival Traits in White-leg Shrimp *(Penaeus vannamei)* Breeding Line"

Table S5. Univariate genetic analyses of the likelihood ratios and the full-sib family effect (*c^2^*± se) for BL2019 breeding lines, significance test at *α* = 0.05 (*χ^2^ _1DF_*).

| Traits | Maximum Log L | | *χ2* Test | *c^2^* ± SE |
| --- | --- | --- | --- | --- |
|  | Model 3 | Model 4 |  |  |
| S1 | 5768.115 | 5777.462 | 18.694 | 0.004 ± 0.005 |
| S2 | -476.013 | -476.142 | 0.258 | 0.003 ± 0.004 |
| S3 | -6452.268 | -6453.577 | 2.618 | 0.006 ± 0.005 |
| S4 | -12344.280 | -12344.839 | 1.118 | 0.005 ± 0.006 |
| S5 | -18021.288 | -18021.311 | 0.046 | 0.001 ± 0.007 |
| S6 | -21096.957 | -21096.711 | 0.492 | 0.000 ± 0.008 |
| S7 | -24353.194 | -24353.023 | 0.342 | 0.000 ± 0.007 |
| S8 | -27927.666 | -27927.591 | 0.150 | 0.000 ± 0.007 |
| S9 | -27927.650 | -27927.591 | 0.118 | 0.000 ± 0.007 |
| S10 | -36742.346 | -36742.328 | 0.036 | 0.000 ± 0.008 |
| S11 | -38357.903 | -38357.896 | 0.014 | 0.000 ± 0.008 |
| S12 | -39270.159 | -39270.156 | 0.006 | 0.000 ± 0.008 |
| SL1 | -42411.363 | -42410.434 | 1.858 | 0.000 ± 0.003 |
| SL2 | -7233.826 | -7233.360 | 0.932 | 0.002 ± 0.004 |
| SL3 | -20866.897 | -20866.782 | 0.230 | 0.001 ± 0.003 |
| SL4 | -31118.638 | -31118.957 | 0.638 | 0.003 ± 0.004 |
| SL5 | -39725.757 | -39726.094 | 0.674 | 0.005 ± 0.005 |
| SL6 | -47308.498 | -47308.692 | 0.388 | 0.003 ± 0.006 |
| SL7 | -53892.299 | -53892.299 | 0.000 | 0.000 ± 0.006 |
| SL8 | -59675.190 | -59675.186 | 0.008 | 0.000 ± 0.007 |
| SL9 | -64908.674 | -64908.673 | 0.002 | 0.000 ± 0.007 |
| SL10 | -69839.754 | -69839.598 | 0.312 | 0.000 ± 0.007 |
| SL11 | -74536.239 | -74536.238 | 0.002 | 0.000 ± 0.008 |
| SL12 | -78907.402 | -78907.402 | 0.000 | 0.000 ± 0.008 |
| SUR | -48671.657 | -48679.731 | 16.148 | 0.128± 0.036 |
| BW | -40784.259 | -40784.453 | 0.388 | 0.030 ± 0.049 |
